# Chronic mycobacteria infection triggers macrophage senescence

**DOI:** 10.1101/2024.11.05.622137

**Authors:** E. Scarpa, D. Oliveri, G. Moschetti, A. Griego, A. Vianello, D. Rondelli, B. Antinori, G. Marchello, P. Santarelli, M. Albanese, S. Franzè, G. Cidonio, L. Manganaro, Nicola I. Lorè, L. Rizzello

**Author notes:** These authors contributed equally to this work (E. Scarpa, D. Oliveri). These authors share the senior authorship (E. Scarpa, L. Rizzello).

## Abstract

Chronic infections with intracellular pathogens such as *Mycobacterium abscessus* (*Mab*) pose significant health challenges due to their capacity to persist within host cells and evade immune responses. This study investigates the cellular responses to chronic Mab infection in macrophages, particularly focusing on cellular senescence. Using an in vitro model of chronic infection in murine alveolar-like macrophages, we found that Mab induces a senescent phenotype characterised by decreased proliferation, altered morphology, DNA damage signalling activation, and upregulation of senescence markers such as p21 and SA-β-galactosidase. Intriguingly, senescent macrophages secreted pro-inflammatory cytokines, consistent with a senescence-associated secretory phenotype (SASP), which promoted secondary senescence in neighbouring uninfected cells. This paracrine transmission of senescence underscores a potentially deleterious effect of *Mab*-induced SASP on tissue microenvironments, fostering a pro-inflammatory niche that may contribute to pathogen persistence. These findings highlight *Mab*-induced senescence as a key factor in chronic infection pathology, suggesting that targeting senescent cells and SASP-related pathways could enhance treatment outcomes in chronic bacterial infections.

## MAIN

The immune system typically clears pathogens that invade the host during a brief, acute phase of infection^1^. However, over millions of years of evolution, pathogenic and opportunistic bacteria have developed sophisticated strategies to survive within intracellular compartments of the host, establishing persistent niches that can result in lifelong chronic infections^2^.

During colonisation, bacteria release a diverse array of virulence factors—molecules manipulating host pathways to facilitate their survival while also damaging the host^3^. Although bacterial mechanisms are often relatively consistent among strains, the host immune response can be complex and unpredictable. In some cases, the response can be self-damaging, leading to long-lasting adverse effects on the host even in the absence of a continued pathogen presence^4,5^. Understanding the long-term effects of chronic infections on the host cells presents considerable challenges and is often overlooked^6^. However, uncovering the molecular mechanisms that enable pathogen immune evasion and chronicity of infection is essential. These insights are likely essential for advancing our understanding of the complex interplay between immune and non-immune cells in modulating immune responses. Moreover, they can shed light on how these cellular interactions shape pathogen dynamics, foster robust immune memory, and help prevent the harmful effects associated with infection and immunopathology^6^.

An intriguing aspect of chronic bacterial infection is its relationship with cellular senescence, a state of irreversible cell cycle arrest that damaged cells often enter in response to prolonged stress^7^. Senescent cells are metabolically active, and remain viable for an extended period of time, being resistant to apoptosis^8^. Also, senescent cells acquire a secretory phenotype, called senescence-associated secretory phenotype (SASP)^9^, characterised by the production of pro-inflammatory cytokines and chemokines (*e*.*g*. IL-1α, IL-1β, and IL6) which promote senescence transmission in a paracrine manner^10^. Cellular senescence occurs in response to different stresses, including DNA damage, oncogenes activation or telomere dysfunction^7^. Intracellular infections can be addressed among the senescence stimuli^11^. Indeed, several viruses were shown to induce cellular senescence in the host by inducing DNA damage or impairing the DNA damage response (DDR)^12,13^. While previous studies have demonstrated that isolated bacterial toxins can similarly induce cellular senescence through DNA damage^14^, the effects of chronic infection involving live bacteria over extended periods remain unexplored. Investigating how ongoing bacterial presence drives cellular senescence is essential for advancing our understanding of long-term host-pathogen dynamics and may reveal novel avenues for therapeutic intervention.

Here, we investigated the effects on the host of a chronic *Mycobacterium abscessus* (*Mab*) infection. *Mab* is an emerging, rapidly growing species of non-tuberculous mycobacterium with intrinsic antibiotic resistance and the ability to elude the immune system ^15–17^. Interestingly, *Mab* can persist intracellularly in macrophages, their natural host, and promote the formation of granuloma structures ^16,18^. We developed an *in vitro* protocol simulating chronic *Mab* infection in murine alveolar-like macrophages. Using this model, we demonstrated that host cells undergo senescence in response to the pathogen’s persistent intracellular presence. Moreover, senescent infected cells secrete an array of SASP factors, transmitting secondary senescence signals to neighbouring uninfected cells.

## RESULTS

### Model of chronic *Mab* infection *in vitro*

To test the hypothesis that chronic intracellular infections caused by *Mycobacterium abscessus* (*Mab*) induce cellular senescence, we used murine alveolar-like macrophages, i.e., the Max Planck Institute (MPI) cells. These self-renewing macrophages recapitulate the innate immune characteristics of resident alveolar macrophages – the natural host for mycobacteria – and represent an exquisite model to frame senescence in the context of host-pathogen interactions^19,20^.

We first established a protocol for chronic infection *in vitro* by exposing cells to a low bacterial load (multiplicity of infection, MOI, equal to 1:0.25) of a *Mab* strain constitutively expressing tdTomato. After an insult of 2 hours, cells exposed to *Mab* were extensively washed to remove residual extracellular bacteria and cultured over 5 days, performing further washes and media change every day, starting from the third-day post-infection (Fig. 1a). This protocol of chronic infection *in vitro* averts premature cellular mortality caused by the overgrowth of extracellular bacteria. It also allows the steady growth of intracellular bacteria (Fig. 1b), leading to 40% ± 17% of infected cells (*Mab*^+^) after 5 days (Fig. 1c). Consistent with the physiological characteristics of alveolar macrophages^21^, unstimulated MPI cells retain the ability to replicate under standard culture conditions^22^. However, a pulse of the EdU-based cellular proliferation analysis revealed a significant reduction upon chronic exposure to *Mab*, with only ∼5% of cells in active replication after 5 days (Fig. 1d,e). Among this small fraction, replicating cells were equally distributed between those infected that harboured the fluorescent bacteria (*Mab*^+^) and those uninfected (*Mab*^−^)(Fig. 1f). Note that doubling the MOI (to 1:0.5) led to an expected increase in the percentage of infection and same reduction in proliferation, but compromised cell viability (Extended Data Fig. 1a,b).

**Figure 1.**
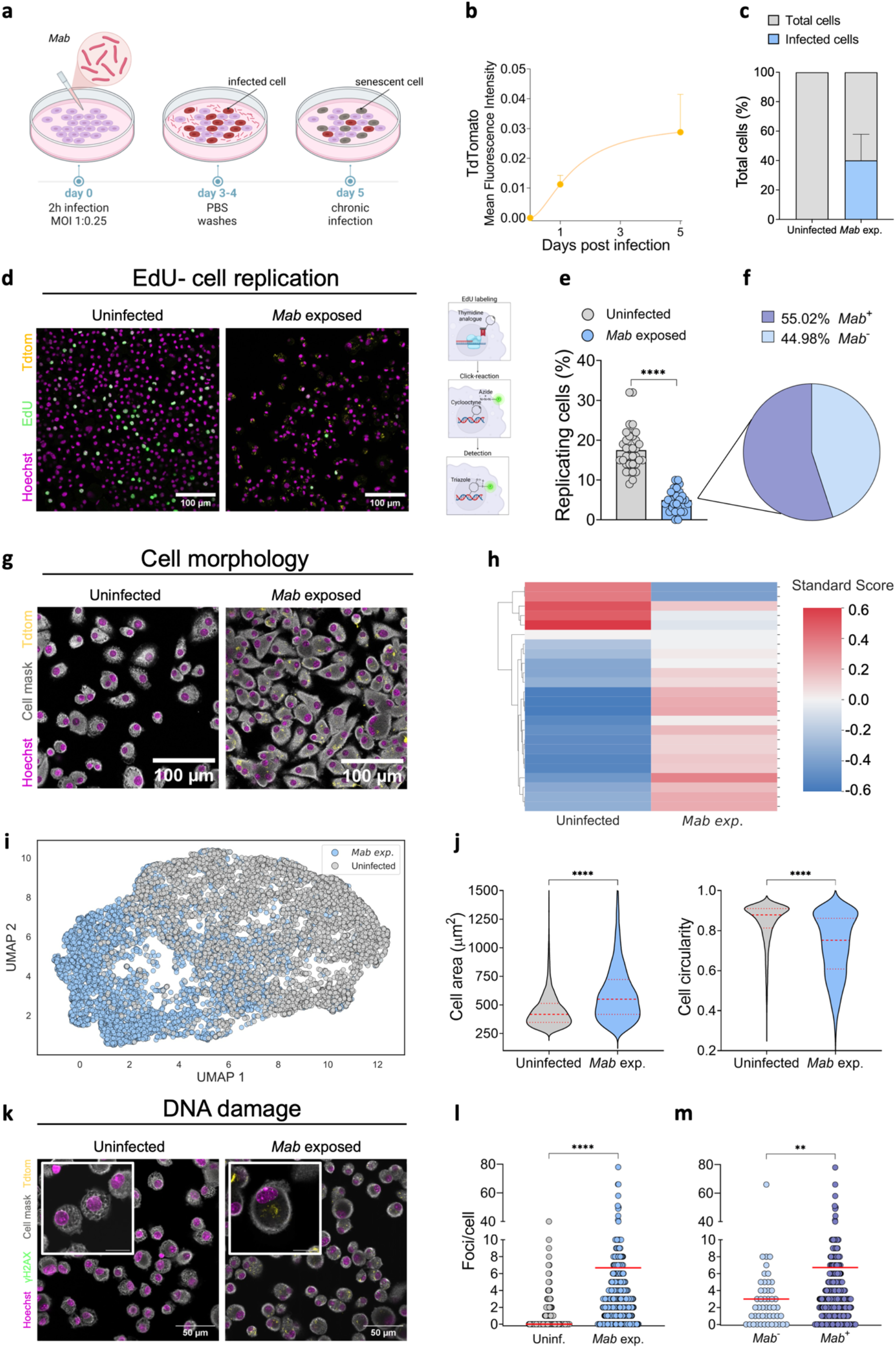
Model of chronic *Mab* infection in macrophages. **a** Schematic representation of the experimental set-up to generate an *in vitro* model of chronic *Mab* infection in alveolar-like macrophages. **b** Quantification of *Mab* intracellular colonisation in alveolarlike macrophages over 5 days exposure. TdTomato mean fluorescence intensity was used as a proxy for *Mab* internalisation. Data showed as mean ± SD. **c** Percentage of *Mab*^+^ cells after 5 days of exposure to the pathogen. Data showed as mean ± SD. **d** Representative confocal microscope images of uninfected or *Mab*-exposed MPI cells stained with EdU. Cell nucleus (Hoechst, magenta), EdU (green) and *Mab* (tdTom, yellow). Scale bar 100μm. Schematic of EdU staining protocol (right). **e** Percentage of replicating cells in uninfected or *Mab*-exposed MPI cells. Data showed as mean ± SD. Non-parametric t-test (Mann-Withney) analysis (**** p<0.0001). **f** Pie chart showing the percentage of *Mab*^+^ or *Mab*^−^ replicating cells in the *Mab* exposed condition. **g** Representative confocal microscope images showing cell morphology in uninfected and *Mab*-exposed cells. Cells (CellMask, grey), Nucleus (Hoechst, magenta) and *Mab* (TdTom, yellow). **h** Heatmap of the standard-score profile in uninfected and *Mab*-exposed cells. The y-axis comprises 24 cellular morphological features (Red = positive modulation, Blue = negative modulation, White = no change). **i** UMAP dimensionality plot of the morphological features in uninfected (grey) and *Mab* exposed (light blue) cells. **j** Violin plot of cell area (left), cell circularity (right) of uninfected and *Mab*-exposed cells. Central dashed line indicates the median, the upper and lower dotted lines indicate the quartiles. Non-parametric t-test (Mann-Withney) analysis (****p<0.0001) Data are from x independent experiments. **k** Representative immunofluorescence images for DNA damage in uninfected and *Mab* exposed cells. Images acquired at 40x magnification, scale bar 100 μm, inset scale bar 25μm. Cells (CellMask, grey), Nucleus (Hoechst, magenta), γH2AX (green) and *Mab* (TdTom, yellow). **l-m** Image analysis for DNA damage signalling activation showing the average number of foci per cell in uninfected or *Mab*-exposed cells (**l**) and the average number of foci per cell in *Mab*^+^ or *Mab*^−^ cells for the *Mab*-exposed ones (**m**). Red line indicates the mean. Non-parametric t-test (Mann-Withney) analysis (** p<0.01; **** p<0.0001).

To better understand the phenotypic effects of long-term exposure to *Mab* on the host cells, we used an image-based approach to quantify cellular morphological changes (Fig. 1g). We employed a high-content multidimensional screening to profile 24 phenotyping features including cell area, perimeter, and mean radius. For each cell and feature, we calculated the standard score, representing the number of standard deviations from the mean of the total population, which included both uninfected and *Mab-*exposed populations^23^. Median scores for each feature were then used to produce a heatmap profile that showed a clear distinction between uninfected and *Mab-*exposed cells (Fig.1h). Further dimension reduction analysis confirmed two clearly resolved clusters of cells having different morphological patterns (Fig. 1i). Several features, including cell area and circularity, were significantly different in the two cellular populations (Fig. 1i,j).

Notably, the discernible phenotypic variances were also evident in the cell nuclei of *Mab-* exposed cells, suggesting a significant change in the nuclear morphological profile (Extended data 1c,d). In this context, we tested whether the changes in nucleus morphology were also associated with the activation of DNA damage signalling (Fig. 1k).

Immunofluorescence staining for γH2AX, a marker for monitoring the activation of DNA damage repair, revealed that cells exposed to *Mab* for 5 days had four times more foci *per* cell compared to uninfected control (*p*<0.0001, Fig. 1l), with *Mab*^+^ cells displaying significantly more foci than *Mab*^−^ cells (Fig. 1m). We hypothesised that such DNA damage signalling activation could be the result of the reactive oxygen species (ROS) produced by host macrophages responding to the chronic intracellular presence of the pathogen. Nevertheless, *Mab-*exposed cells did not produce any ROS (Extended Data Fig. 1e), most likely as a part of an immune evasion mechanism of the pathogen. Thus, the mechanism inducing DNA damage signalling activation remains elusive^24^.

The findings collectively establish a model of chronic *Mab* infection in macrophages and demonstrate that chronically infected cells reduce the proliferation rate, change cellular and nuclear morphology, and activate the DNA damage signalling. All these features are classically associated with the onset of cellular senescence^25^.

### Chronic *Mab* infection induces cellular senescence in macrophages

To confirm that the long-term presence of intracellular *Mab* is the trigger for senescence, rather than a persistent inflammatory stimulation, we examined the expression of a panel of senescence-related cellular markers after exposure to *Mab* or lipopolysaccharide (LPS) for up to 5 days (Fig. 2a).

**Figure 2.**
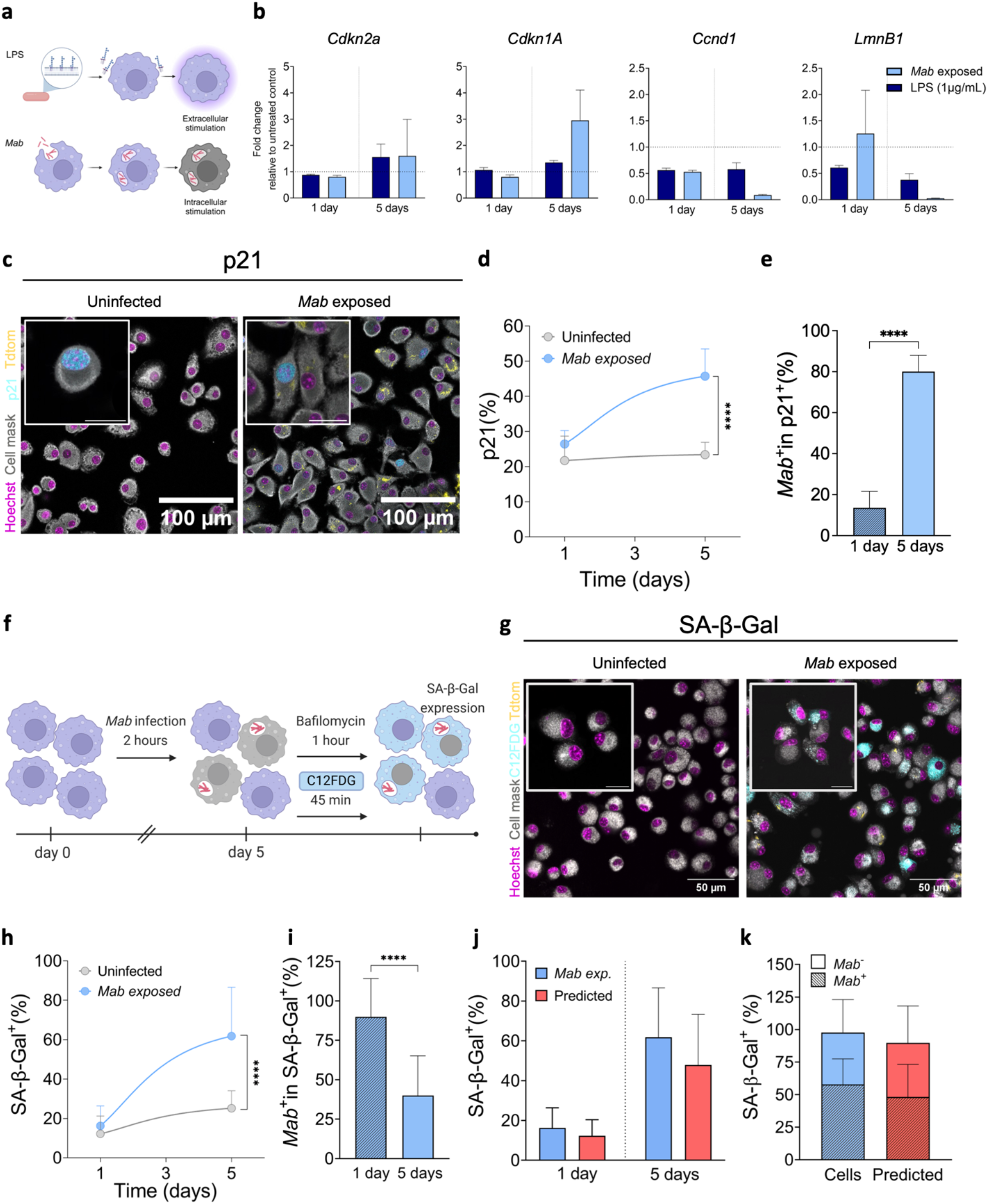
Chronic Mab infection induces cellular senescence in macrophages. **a** Schematic representation of MPI activation upon LPS stimulation (upper panel) or *Mab* exposure. **b** Differential expression of senescence-associated genes for LPS treated (1μg/mL) or *Mab* exposed cells, in acute or chronic condition. Data normalised on *36b4* housekeeping gene and untreated control. Normalised data expressed as fold change (2^-ΔΔCt^). Data showed as mean ± SD. **c** Representative immunofluorescence images for p21 expression. Images acquired at 40x magnification, scale bar 100 μm, inset scale bar 2 5μm. Cells (CellMask, grey), Nucleus (Hoechst, magenta), p21 (light blue) and *Mab* (TdTom, yellow). **d** Time-dependent variation of the percentage of p21 positive cells. Non-parametric t-test (Mann-Withney) analysis (**** p<0.0001). **e** Percentage of *Mab*^+^ cells among the p21 positive group. Data showed as mean ± SD. Non-parametric t-test (Mann-Withney) analysis (*p<0.05). **f** Schematic of SA-β-gal (*i*.*e*., C12FDG) staining protocol upon chronic *Mab* infection. **g** Representative immunofluorescence images for SA-β-Gal expression. Images acquired at 40x magnification, scale bar 100μm, inset scale bar 25μm. Cells (CellMask, grey), Nucleus (Hoechst, magenta), C12FDG (light blue) and *Mab* (TdTom, yellow). **h** Time-dependent variation for the percentage of SA-β-Gal expressing cells. Non-parametric t-test (Mann-Withney) analysis (****p<0.0001). **i** Percentage of *Mab*^+^ cells among the SA-β-Gal expressing cells. Data showed as mean ± SD. Non-parametric t-test (Mann-Withney) analysis (****p<0.0001). **j,k** Machine learning-based predictive analysis for SA-β-Gal positive cells (**j**) and for Mab^+^ or *Mab*^−^ cells among the SA-β-Gal positive group (**k**) compared to *in vitro* experiments.

Transcriptomic analyses over senescence-associated genes revealed no significant alterations after 24 hours of stimulation, irrespective on the origin of the insult (either LPS or *Mab*). However, cells exposed to *Mab* for 5 days exhibited upregulation of *Cdkn2a* (p16^ink4a^, hereafter p16) and *Cdkn1a* (p21^Cip1^, hereafter p21), two genes regulating cell-cycle progression, alongside the downregulation of *Ccnd1* and *LmnB1*, also involved in controlling cell-cycle nuclear structure, respectively. This pattern was not observed with LPS incubation, where genes involved in the cell-cycle remained unaltered (Fig. 2b), confirming the distinct feature of *Mab*-triggered senescence as compared to a classical LPS-induced activation.

The expression of p21 was further validated by immunofluorescence (Fig. 2c), revealing a significant increase in the proportion of p21^+^ cells among the *Mab*-exposed cells (45.7% ± 7.7%) compared to uninfected controls, which maintained steady levels of expression over time (*p* < 0.0001) (Fig. 2d).

Among cells exposed to the pathogen, the proportion of p21^+^ cells within the *Mab*-harbouring population (Mab^+^) was 25.0% ± 2.6% after 1 day, increasing significantly to 80.0% ± 7.9% by day 5 (*p* < 0.0001) (Fig. 2e).

To further confirm the senescence phenotype associated with chronic *Mab* infection, we assessed the expression of senescence-associated β-galactosidase (SA-β-gal) following chronic exposure to the pathogen. To reduce intrinsic basal levels of SA-β-gal in macrophages^22^, we implemented a protocol incorporating pre-treatment with bafilomycin to inhibit endogenous β-galactosidase activity (Fig. 2f). Cells chronically exposed to *Mab* exhibited intense cytoplasmic staining, consistent with the lysosomal localisation of SA-β-gal (Fig. 2g). A time-course analysis revealed a progressive significant increase in the proportion of *Mab*-exposed cells positive for SA-β-gal (∼60% positive cells at day 5, *p* < 0.0001) (Fig. 2h). Among the SA-β-gal^+^ population, approximately 80% of cells were also Mab^+^ at day 1. However, a significant reduction in the number of *Mab*^+^ SA-β-gal^+^ cells was observed at day 5 (*p* < 0.0001), indicating that senescence also occurred in cells devoid of *Mab* (Fig. 2i).

To confirm the onset of cellular senescence in cells exposed to *Mab*, we utilised a recently developed machine-learning classifier based on nuclear morphology features^26,27^. Using a transfer learning approach, we retrained the algorithm with nuclear features from healthy MPI cells as a negative control. Quality parameters validated the newly developed tree-based classifier, which was subsequently applied to predict cellular senescence in our chronic infection model. The percentage of SA-β-gal^+^ cells closely matched the classifier predictions in *Mab*-exposed cells (Fig. 2j) as well as in *Mab*^+^ and *Mab*^−^ populations (Fig. 2k). Collectively, these findings proved the hypothesis that chronic *Mab* infection induces cellular senescence in macrophages.

### Paracrine transmission of senescence from infected cells

After demonstrating that *Mab* chronic infection pushes macrophages towards a senescent phenotype, we assessed the paracrine effects on the surrounding uninfected cells. Cellular senescence can be transmitted to neighbouring cells via paracrine signalling, a process termed secondary senescence, which has been demonstrated in various pathological contexts, including cancer progression and tissue fibrosis^28^.

We characterised the distribution and spread of cellular senescence from *Mab*^+^ cells to *Mab*^−^ bystanders to probe whether senescence triggered by chronic infection can be transmitted to neighbouring (non-senescent) cells. Image analysis was performed to assess the spatial distribution of SA-β-gal^+^ cells in relation to infected cells within a 60 or 120 μm radius (and locating one infected cell at the centre of the circle), corresponding to distances of 2 or 4 cells, respectively (Fig. 3a). There is evidence that the close proximity of an infected cell is characterised by a high prevalence of bystander uninfected SA-β-gal^+^ cells (85.7% ± 14.3%) suggesting that senescence is closely associated with infected cell clusters (Fig. 3b).

**Figure 3.**
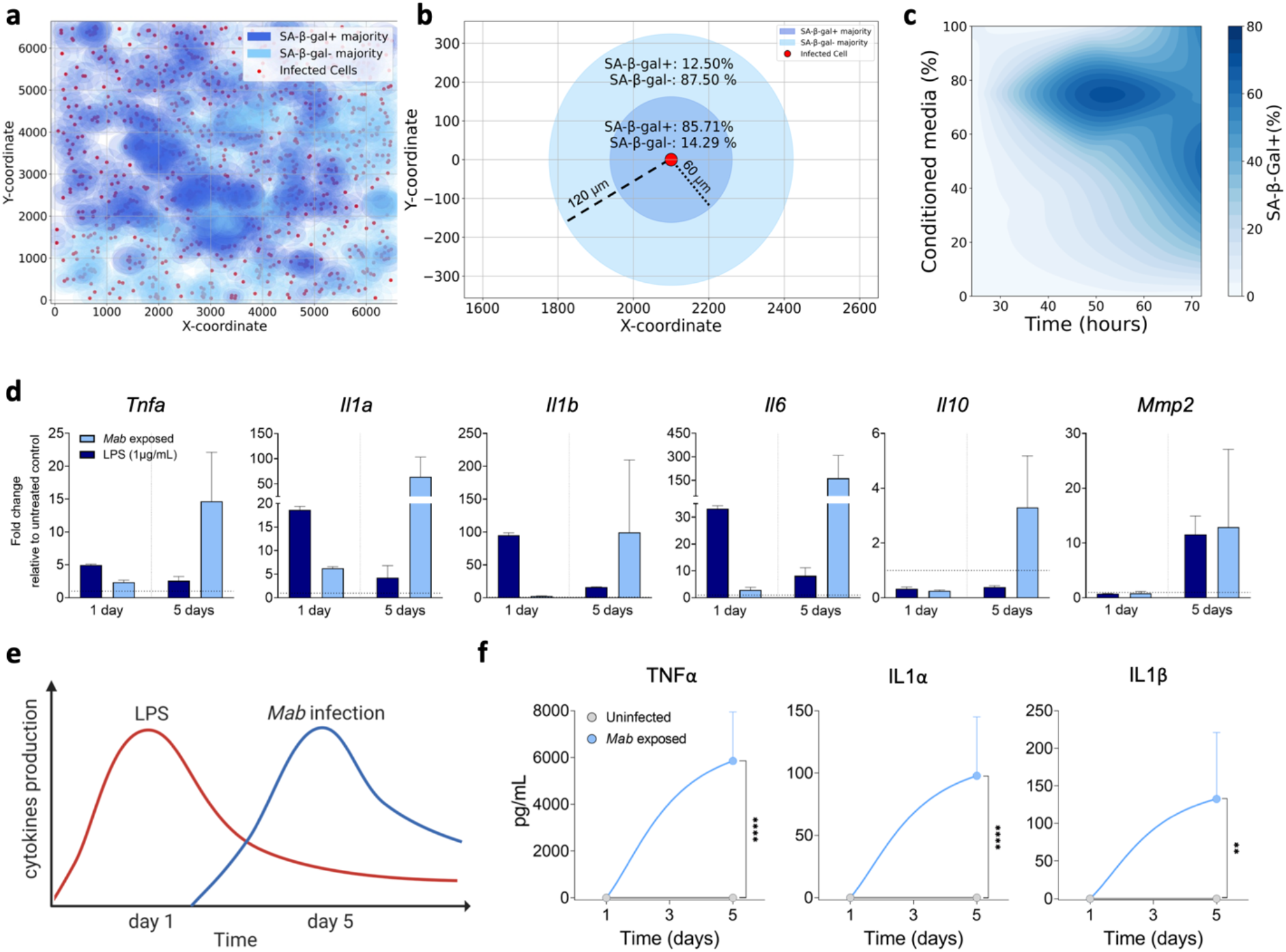
Chronic Mab infected macrophages produce SASP and transmit cellular senescence. **a,b** Neighbourhood analysis for senescence transmission. **a** Visual representation of *Mab*^+^ coordinates (red dots) and surrounding prevalence of SA-β-Gal^+^ (dark blue) or SA-β-Gal^−^ cells (light blue). **b** Quantitative analysis showing the percentage of SA-β-Gal^+^ or SA-β-Gal^−^ cells within a range of 60 or 120 μm. **c** Contour plot showing the percentage of SA-β-Gal positive cells over time upon incubation with different dilutions of conditioned media isolated from *Mab-*exposed cells. **d** Differential expression of SASP-associated genes in LPS treated or *Mab* exposed cells, in acute or chronic condition. Data normalised on *36b4* housekeeping gene and untreated control. Normalised data expressed as fold change (2^-ΔΔCt^). Data showed as mean ± SD. **e** Schematic representation of different dynamics in cytokines production for extracellular stimulus, LPS, and intracellular chronic stressor, *Mab*. **f** ELISA results for SASP-related protein production (TNFα, ILα, ILβ) control vs *Mab* exposed cells, in acute or chronic condition.

To confirm the paracrine induction of senescence, we collected conditioned media from cells exposed to *Mab* for 5 days. Upon filtering it, to eliminate any remaining extracellular bacteria, the conditioned media was added at varying ratios to healthy uninfected cells, which were then screened by confocal microscopy and imaging analyses. Cells exposed to conditioned media exhibited increasing levels of SA-β-gal in a dose-dependent manner (growing ratio of conditioned/fresh media mixture), reaching ∼80% positivity as early as 48 hours after incubation with 75% conditioned media (Fig.3c). This suggested that factors secreted by infected cells (most likely the SASP, investigated next) contribute to the senescence phenotype. Notably, cells exposed to *Mab* expressed high levels of pro-inflammatory genes associated with the SASP only after chronic exposure to the pathogen. In contrast, incubation with LPS for up to 5 days induced upregulation of these genes only during the first 24 hours, except for metalloprotease-2, which returned to near baseline levels thereafter (Fig. 3d). Gene expression quantification was corroborated by a significant increase in secretion of pro-inflammatory cytokines such as TNFα, IL-1α, and IL-1β after 5 days of *Mab* exposure (Fig. 3e). Overall, the release of pro-inflammatory molecules follows two distinct patterns depending on the stimulus: while LPS triggers an acute (fast) response that progressively declines over time (thus unrelated to senescence), *Mab* exposure triggers a delayed and persistent increase of pro-inflammatory molecules coinciding with senescence induction (Fig. 3f). The evidence showed here indicate that chronically infected senescent cells can induce senescence in neighbouring uninfected cells via SASP factor secretion.

## DISCUSSION

Cellular senescence is a biological state of irreversible growth arrest and metabolic alterations in response to different triggers and chronic insults that promote, or exacerbate, different pathological conditions^25^. Senescent cells are also characterised by a pro-inflammatory secretome, and by increased resistance to apoptosis. Although the onset of cellular senescence may represent an innate mechanism to prevent the unrestricted growth of damaged cells^8^, it may also create a permissive environment for persistent infections.

Cellular senescence has been reported to occur during viral infections^12^ and as an indirect consequence of bacterial colonisation. Bacteria, both pathogenic and commensal, can promote senescence either through secreted metabolites, like deoxycholic acid and butyrate, or by persistently activating pattern recognition receptors that induce senescence pathways^29–31^. Alternatively, bacteria can directly induce cellular senescence by secreting genotoxins that cause DNA damage in the host cells^32,33^. In particular, *Salmonella enterica* secretes the typhoid toxin as part of its virulence strategy, which triggers a senescent-like phenotype in infected cells and leads to the secretion of SASP-related factors that transmit senescence to neighbouring cells^14^.

Despite these insights, the role of senescence in chronic infections caused by intracellular bacteria remains poorly understood. In this study, we developed an *in vitro* model of chronic *Mab* infection to assess the long-term effects of living intracellular pathogens on the host. *Mab* is an emerging, rapidly growing species of non-tuberculous mycobacterium with intrinsic antibiotic resistance abilities^15–17^ Using our model, we demonstrated that chronic exposure to *Mab* induces cellular senescence in macrophages, with notable paracrine effects on uninfected bystander cells.

Cells exposed to *Mab* for 5 days displayed a series of classical markers associated to cellular senescence^25^. These included the morphological alterations at both the cellular and nuclear level^23,26^ reduced proliferation rate, impaired control of the cell-cycle^8^, and the activation of the DNA damage repair signalling^34^. Previous reported data demonstrated that mycobacteria, particularly *Mycobacterium tuberculosis* (*Mtb*), *can* promote DNA damage and impair the repair pathways to create a nutrient-rich replicative niche^35^. Although the molecular mechanism that activates the DNA damage signalling, upon *Mab chronic* infection, is yet to be characterised. it is unlikely to be ROS-dependent, as previously demonstrated with *Mtb*^*36*^. The absence of oxidative stress suggests an adaptive strategy of *Mab* to persist intracellularly through immune evasion.

Our results highlight that *Mab*-induced senescence is associated with complex host-pathogen interactions extending beyond traditional immune activation. The upregulation of cell-cycle inhibitors like p21, and increased SA-β-gal activity in *Mab-*exposed cells, underscores that the chronic intracellular presence of the pathogen disrupts cell-cycle regulation, a hallmark of cellular aging^7^. Notably, the up-regulation of senescent-associated genes was absent in cells exposed to LPS, underscoring that *Mab*-induced senescence is not merely a response to generalised inflammation but rather a distinct consequence of intracellular infection. It remains to be elucidated whether the onset of cellular senescence is caused exclusively by *Mab* infection, or it is a recurrent mechanism that pathogens exploit to perpetuate the infection^14^. Further, our analysis reveals significant differences in SASP-related transcriptomic profiles between *Mab* and LPS exposure. While LPS induces an acute, transient inflammatory response, Mab infection sustains a chronic inflammatory state characterized by prolonged secretion of TNFα, IL-1α, and IL-1β. This ongoing inflammatory signaling supports the hypothesis that chronic Mab infection establishes a unique immune microenvironment, which deviates significantly from acute infection models and may contribute to a persistent pro-inflammatory niche within infected tissues.

One of the most intriguing findings of our study is the role of SASP in spreading senescence to neighbouring, uninfected cells, demonstrating a paracrine mechanism of senescence transmission. The observation that uninfected bystander cells acquire senescence markers, including SA-β-gal, has substantial implications for chronic infections, as clusters of senescent cells could foster a pro-inflammatory environment that impairs host defenses, potentially facilitating the persistence and spread of infection within the host tissue.

In conclusion, our study establishes that chronic *Mab* infection induces cellular senescence in macrophages through direct infection-related stress and secondary SASP-mediated effects. These findings highlight two promising therapeutic targets. First, the selective elimination of both infected and bystander senescent cells using senolytic compounds could serve as a host-directed therapy to complement standard antimicrobial treatment, potentially reducing bacterial load and alleviating SASP-induced inflammation^37,38^. Given that senescent cells may serve as reservoirs for chronic infection, senolytics could help eradicate persistent infection niches and reduce the risk of relapse. Second, targeting the SASP pathway to mitigate *Mab*-induced senescence may slow disease progression in chronic Mab infections. Future studies should explore the molecular mechanisms of *Mab*-induced senescence *in vivo*. Additionally, our model of chronic infection offers a valuable platform for evaluating novel therapeutic strategies, including SASP inhibitors and anti-senescent treatments, to improve outcomes for patients suffering from chronic *Mab* infections.

## ACKNOWLEDGMENTS

The authors acknowledge the scientific and technical assistance of Dr Alessandra Fasciani and Dr Chiara Cordiglieri of the INGM Imaging Facility, and Dr Mariacristina Crosti of the Sorting Facility at INGM. We also thank Prof. Marina Freuderberg (BIOOS Centre for Biological Signalling Studies, University 456 of Freiburg) for kindly providing MPI-2 cells.

This work was funded by grants to: E.S. from Fondazione Cariplo (grant number 2022-0438) and Italian Ministry for University and Research (through the Seal of Excellence program, grant number SOE_0000114); L.R from ERC-2019-STG (grant number 850936), the Fondazione Cariplo (grant number 2019-4278), and the Italian Ministry for University and Research (through the PRIN program, grant number 20205B2HZE, the PRIN2022 grant number 2022PL44LM, and the PRIN2022-PNRR grant number P2022CC2N8_002); S.F. from the European Union — NextGenerationEU (PNRR M4C2-Investimento 1.4-CN00000041-23 PNRR_CN3RNA_SPOKE8).

## AUTHOR CONTRIBUTIONS

S.E., O.D., and R.L. designed the study, while S.E., O.D., M.G., G.A., V.A., R.D., A.B., and S.P. developed the methodology. Data curation was carried out by S.E., O.D., M.G., G.A., V.A., R.D., and A.B., with validation efforts contributed by S.E., O.D., M.G., G.A., V.A., R.D., A.B., and S.P. The original draft was written by S.E., O.D., and R.L., and the review and editing process involved M.G., G.A., V.A., R.D., A.B., S.P., A.M., F.S., C.G., M.L. and L.I.N. Supervision was provided by S.E. and R.L., who also acquired the necessary funding for the study.

### Lead contacts

Further information and requests for resources and reagents should be directed to and will be fulfilled by the Lead Contacts, Edoardo Scarpa or Loris Rizzello

## MATERIAL AND METHODS

### Materials Availability

Plasmids and bacterial strains generated in this study are available from the Lead Contact with a completed Materials Transfer Agreement.

### Murine alveolar-like macrophages

Max Plank Institute (MPI) cells (alveolar-like macrophages) at passage below 40, were cultured in RPMI 1640 medium (Thermo Scientific) supplemented with 10% heat-inactivated Fetal Bovine Serum (Euroclone), 1% Glutamine (Euroclone), 1% Penicillin and Streptomycin (Euroclone), 20 mM Hepes (Euroclone) and 0,6 µg/mL of GM-CSF (Peprotech). Cell cultures were maintained in the incubator at 37°C with 5% CO_2_. Cells were split twice a week and plated at 2 × 10^5^ cells/mL to avoid confluence. For chronic LPS stimulation, media containing LPS (1μg/mL) was replaced daily, for up to 5 days.

### Mycobacterium abscessus Tdtomato fluorescent reporter strain

A reporter strain of *Mycobacterium abscessus* (*Mab)* constitutively expressing the fluorescent reporter TdTomato was used in the experiments. This *Mab* strain carries an integrative plasmid designed to constitutively express the *tdTomato*-L5 gene (Addgene #140994) under the control of a strong promoter, HSP65. *Mab* strain was cultured at 37 °C in Middlebrook 7H9 broth base supplemented with Middlebrook OADC (oleic acid, albumin, dextrose, and catalase) growth supplement (Fisher Scientific), Tween 80 (Sigma-Aldrich) and 0.5% of glycerol. For the infection, MPI cells were plated at 2×10^5^ cells/mL one day in advance, then infected at the indicated MOI (1:0.25 or 1:0.5) for 2 hours, finally washed with PBS three times and incubated at 37°C 5%CO_2_. To obtain a model for chronic infection and to avoid cell death due to the extracellular growth of *Mab*, the medium was replaced daily from day 3 post-infection.

### EdU staining

To assess cell proliferation upon chronic infection, EdU fluorescent staining was performed using Click-it EdU cell proliferation kit (Thermo Scientific). Cells were labelled with EdU (5-ethynyl-2’
s-deoxyuridine) (10 μM) for 1 hour then fixed with 4% (v/v) paraformaldehyde (PFA) in PBS for 15 minutes. Cell permeabilization was performed using 0.5% (v/v) Triton X-100 in PBS. For detection, Click-it reaction, containing AlexaFluor-488 fluorophore, was prepared following manufacturer instructions. After 30 minutes of incubation, protected from light, cells were washed with 3% (v/v) Bovine serum albumin (BSA) in PBS. Additional staining was performed for nuclei visualisation using Hoechst 1:1000 in PBS (Cell signalling) for 15 minutes. Cells were maintained in PBS at 4°C until the analysis.

### Immunofluorescence

Cells were fixed with 4% (v/v) PFA for 20 minutes, washed twice in PBS, permeabilised with 0.5% (v/v) Triton™ X-100 for 10 minutes and blocked with 5% (v/v) BSA for 1 hour at room temperature. Cells were incubated with primary antibody anti-phospho-histone H2A.X (γH2AX) (#2577; Cell Signaling. 1:200 in 5% BSA in PBS), for 1 hour, or Waf1/Cip1/CDKN1A p21 (sc-6246, Santa Cruz. 1:200 in 1% BSA in PBS) for 3 hours. After 3 washes with PBS, cells were incubated with secondary antibody Alexa Fluor® 488-Goat anti-mouse IgG (Thermo Scientific. 1:500 in 1% BSA in PBS) for 1 hour. Finally, Hoechst (Cell Signalling. 1:500 in PBS) and CellMask™ Deep Red Plasma membrane (Thermo Scientific. 1:2000 in PBS) were added for 15 minutes at room temperature. Cells were then washed with PBS and stored at 4°C until analysis.

### C12 FDG staining

Cells were stained with Hoechst (Cell Signalling. 1:500 in RPMI) and CellMask™ Deep Red Plasma membrane (Thermo Scientific. 1:2000 in RPMI) for 20 minutes at 37°C. After a wash in PBS, cells were pre-treated with Bafilomycin A1 (HY-100558, MedChemExpress) 100 nM for 1 hour, then stained with C12FDG (ab273642; Abcam) 33μM for 45 minutes at 37°C. Cells were then fixed with 4% (v/v) PFA for 20 minutes and stored in PBS at 4°C until analysis.

### High-resolution confocal microscopy

Fluorescence stainings were visualised with Leica STELLARIS 8 Confocal Microscopy using LasX software (Leica). For every sample, 10-60 fields of view with 20x or 40x magnification objective were acquired, for a total of up to 4000 cells per condition. Quantitative microscopy analysis was performed using NIS-Elements Advanced Research (Nikon) software or CellProfiler 4.2.7. The morphological features analysed were extracted using CellProfiler 4.2.7 and are listed here: *MedianRadius, Eccentricity, Solidity, Max Feret Diameter, Maximum Radius, Equivalent Diameter, Major Axis Length, Bounding Box Maximum X, Mean Radius, Bounding Box Minimum_X, Min Feret Diameter, Perimeter, Area, Spatial Moment_0_0, Minor Axis Length, Form Factor, Convex Area, Bounding Box Area, Extent, Spatial Moment_0_1, Normalized Moment_2_1, Bounding Box Maximum_Y, Bounding Box Minimum_Y, InertiaTensor Eigen values_1*.

### Conditioned media

Conditioned media was collected on the fifth-day post-infection (MOI 1:0.25) and filtered using a 0.22 µm syringe filter. Several dilutions were performed adding RPMI to the filtered media and supplemented with 0,6 µg/mL of GM-CSF. Healthy MPI were then exposed to different concentrations of conditioned media for up to 72 hours, changing the media daily. Cells were then stained with C12FGD and analysed through confocal microscopy.

### Quantitative PCR with reverse transcription

Total RNA was extracted from murine following the manufacturer’s instructions on Monarch® Total RNA Miniprep Kit (T2010S, New England Biolabs). Extracted RNA was quantified with Thermo Scientific™ NanoDrop™ One/OneC Microvolume UV-Vis Spectrophotometer. The reverse transcription was performed with ProtoScript® II First Strand cDNA Synthesis Kit (E6560S, New England Biolabs) using the optimised Random Primer Mix provided by the manufacturer. cDNA produced was diluted 1:4 in nuclease-free water and 5µL were added in each well of a 96-well plate where the forward and reverse primers for the gene of interest and SensiFAST™ SYBR® Lo-ROX Kit (BIO-94020, Meridian Bioscience) were mixed. The RT-qPCR was performed using a Quant Studio Real Time PCR System (Thermo Scientific), all the samples were measured in technical triplicate and the curves of amplification were analysed with Design and Analysis Software Version 2.6 (Thermo Scientific). Expression data were normalised to *36b4* gene. Fold change was calculated using 2^−ΔΔCt^ method. All primer sequences were reported in the Supplementary Tab.

### ELISA

The supernatants derived from uninfected or *Mab*-exposed cells were harvested and stored at -80°C. Assessment of TNFα, IL1α and IL1β production was performed using a commercially available kit (Thermo Scientific), following the manufacturer’s protocols. (TNFα: 88-7324-22, IL1α: 88-5019-22, IL1β: 88-7013-22).

### ROS assay

To detect the production of reactive oxygen species (ROS), H2DCFDA (2’,7’-Dichlorodihydrofluorescein diacetate) (Sigma-Aldrich) was added to the medium at a final concentration of 50 μM for 30 minutes. In the presence of ROS, the substrate is converted into dichlorofluorescein, which emits fluorescence at 522 nm. Fluorescence was measured using Spark® multimode microplate reader (Tecan). As positive control, cells were treated with 50 mM of tert-Butyl hydroperoxide (t-BHP) (Sigma-Aldrich-) for 6 hours.

### Classifier to identify senescent cells

For Classification tree (CT) algorithm python version 3.7.7 was used. The following packages were also utilised: scikit-learn and derived packages, pandas, numpy, matplotlib. pyplot, seaborn, and csv. For the cell and nucelar morphological analysis the public software CellProfiler (version 4.2.4) was used. The training set used and the structure of the CT were obtained from a published dataset^26^. However, the dataset used to train the negative recognition (non-senescent cells) was expanded using data derived from uninfected MPI cells cultivated for 24 hours. For classification tree (CT)-based classifiers, preliminary classification trees were built using scikit-learn, providing 30% of the training set as test size. After assessing the initial accuracy, AUC, and ROC curves, cost complexity pruning was performed to avoid overfitting.

### Statistical analysis

GraphPad Prism Software Version 8 was used for statistical analyses. Two-tailed Student’s t-tests to compare two experimental groups, otherwise, for more groups one-way ANOVA with and Tukey’s multiple comparison test was performed. Non-parametric analysis Kruskal-Wallis or Kolmogorov-Smirnov were performed for not normally distributed data. Data handling was performed using Spyder 5.5.1 on Anaconda Navigator 2.5.2.

#### Primers list

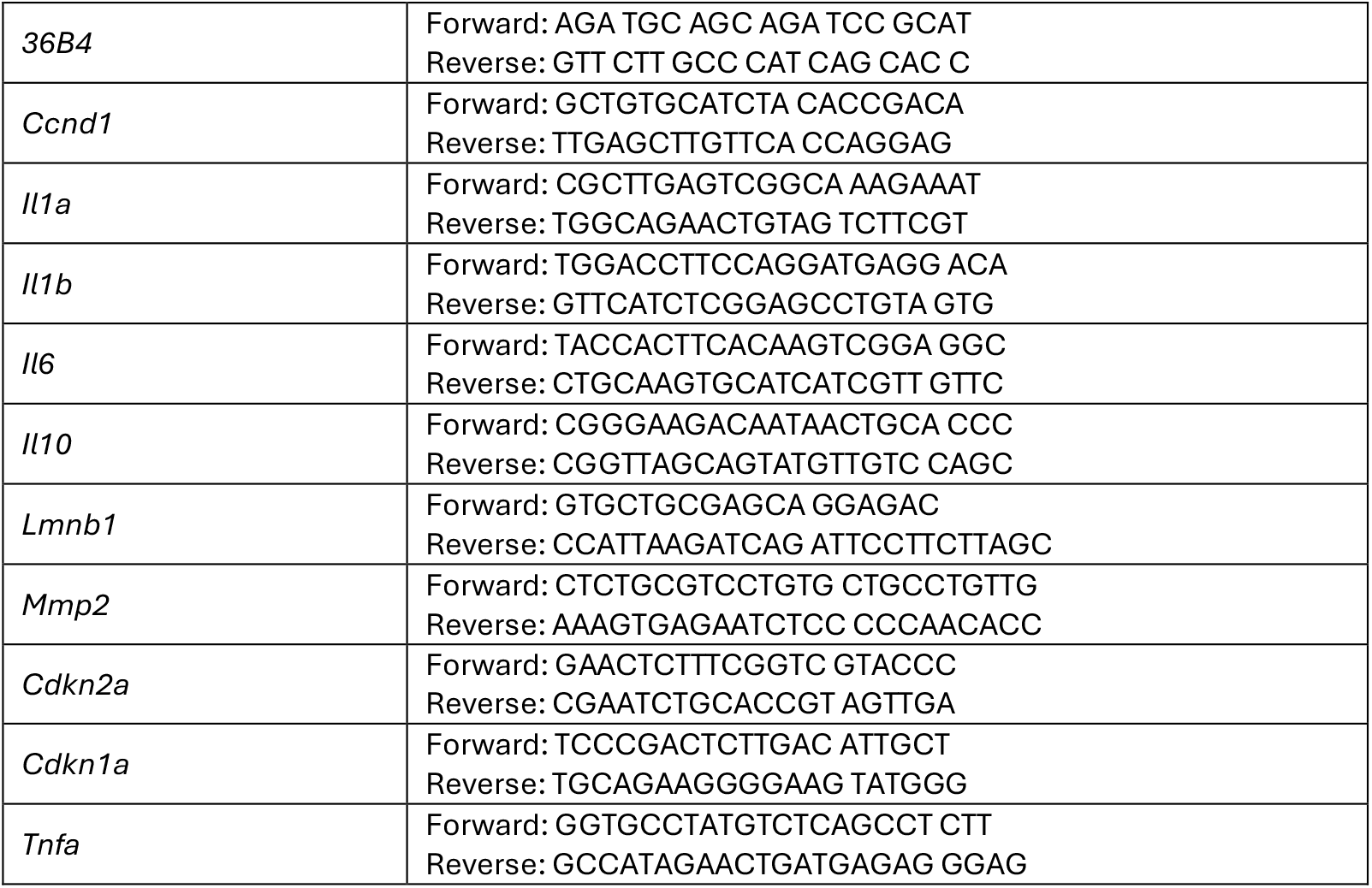

---

**Supplementary figure.**
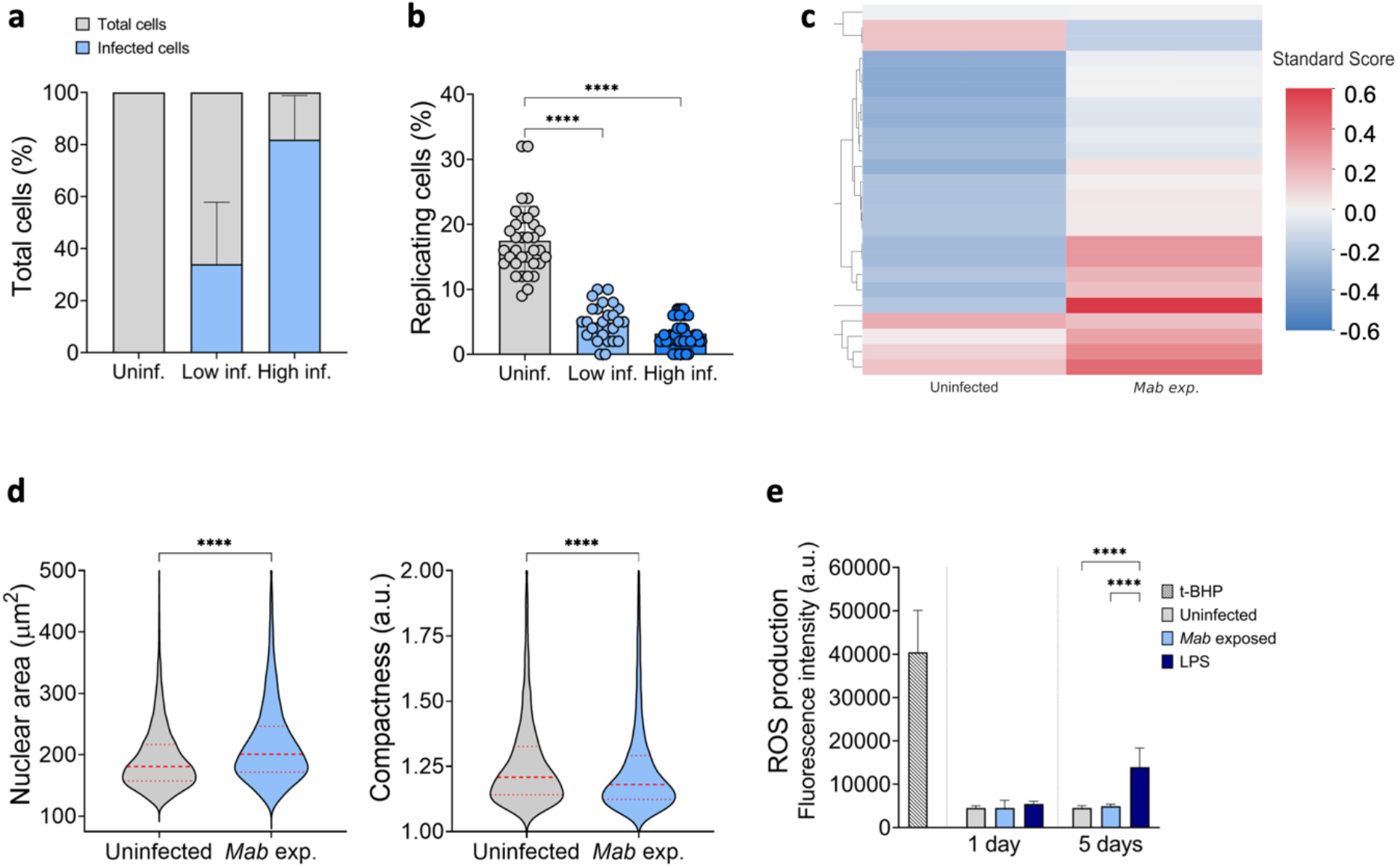
**a** Percentage of *Mab*^+^ cells upon chronic infection at different MOI: 1:0.25 (low infection) and 1:0.5 (high infection). **b** Image analysis performed for the percentage of replicating cells in uninfected, low, and high infected cells. Data showed as mean ± SD. Non-parametric t-test (Mann-Withney) analysis (****p<0.0001) **c** Z-score profile heatmaps for uninfected and *Mab* exposed cells. Y-axis comprises 24 nuclear morphological features (Red = positive modulation, Blue = negative modulation, White = no change) **d** Image analysis of nuclear area (left), nuclear compactness (right). Central dashed line indicates the median, and the upper and lower dotted lines indicate the quartiles. Statistic non-parametric t-test was performed for significance (**** p<0.0001). **e** Fluorescence assay for ROS production in uninfected, *Mab* exposed, LPS treated cells, at 1 and 5 days timepoints. Data showed as mean ± SD. One-way ANOVA comparison analysis (****p<0.0001).

